# Seven inducible promoters for *Zymomonas mobilis*

**DOI:** 10.64898/2026.03.17.712268

**Authors:** Gerrich Behrendt

## Abstract

*Zymomonas mobilis* is an ethanologenic Alphaproteobacterium with many interesting characteristics for fundamental research and applied microbial engineering. Although genetic engineering has been established for *Z. mobilis* since the 1980s, a rich set of inducible transcriptional regulators is still unavailable. In this work, seven different chemically inducible promoters have been systematically tested for their functionality in *Z. mobilis*. In particular, for the first time, NahR-P_salTTC_, VanR^AM^-P_vanCC_, CinR^AM^-P_cin_ and LuxR-P_luxB_ have been characterized in *Z. mobilis*, alongside the commonly used regulator-promoter pairs TetR-P_tet_ and LacI-P_lacT7A1_O3O4_, and the less commonly used XylS-P_m_. All promoters investigated in this work are compatible with the Golden Gate modular cloning framework Zymo-Parts. Characterization was carried out with a shuttle vector backbone based on pZMO7, which has so far been rarely used for applications in *Z. mobilis* but seems to be completely stable without selection and generates high and uniform levels of expression. From the experimental results presented, it can be concluded that VanR^AM^-P_vanCC_ and CinR^AM^-P_cin_ are particularly promising for broad use in the *Z. mobilis* community.

**Graphical abstract:** 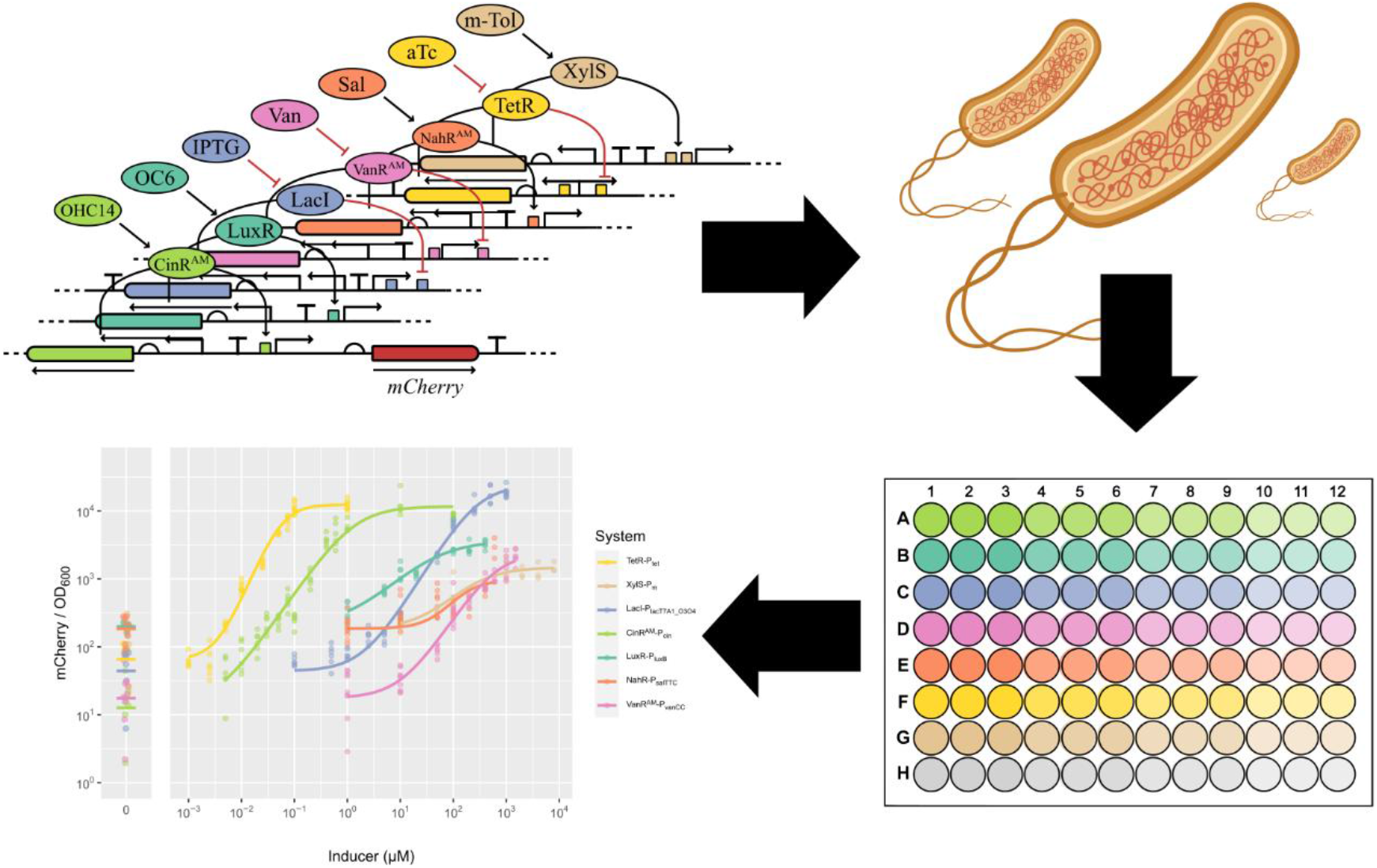

## Introduction

*Zymomonas mobilis* is an ethanologenic Alphaproteobacterium with many interesting characteristics for fundamental research and applied microbial engineering. It has been proposed as a model system for naturally genome-reduced Alphaproteobacteria [1], has a remarkably high hopanoid content in its membrane [2] that allows for growth and survival at high ethanol concentrations [3], divides asymmetrically [4], is polyploid [5] and has an “uncoupled respiration” phenotype [6]. Furthermore, it has some of the highest specific glucose consumption rates for bacteria [7] which makes it an interesting candidate for industrial microbial catalysis. Many heterologous products have already been shown to be producible in *Z. mobilis* to varying degrees of success [8–10].

Though there is ongoing research interest for *Z. mobilis* and ongoing genetic tool development [11–14], most researchers either rely on TetR-P_tet_ or a version of LacI-P_lac_. Both of these transcription regulators work well in *Z. mobilis* [11], but both also have drawbacks. Induction of TetR-regulated systems typically uses tetracycline derivatives (e.g., anhydrotetracycline, aTc), which can still impair growth, albeit less than tetracycline itself [15]. Isopropyl-β-d-thiogalactopyranosid (IPTG), which is commonly used for LacI-regulated transcription is relatively expensive and thus increases costs and can limit large-scale applications. Furthermore, when screened for crosstalk in *E. coli*, the inducer for TetR had comparably strong activity with the LacI-regulated promoter [16], which most likely holds true for applications in *Z. mobilis*. Even without the need to overcome the drawbacks of TetR-P_tet_ and LacI-P_lac_, having more choices for inducible expression systems in *Z. mobilis* would be beneficial in its own right.

In *E. coli*, a directed evolution strategy was developed by Meyer et al. 2019 to create optimal inducible transcription regulators [16]. Here, the researchers started with naturally occurring and synthetic regulator-promoter circuits and selected for optimal performance. They were looking at lower basal activity, high dynamic range, increased sensitivity and low crosstalk. This yielded many different *E. coli* strains that either carry the transcription regulation circuits on a plasmid or have them integrated into the chromosome. Later on, many of the expression systems optimized for function in *E. coli* and systems of different origin were systematically tested in different Proteobacteria and integrated into a HiFi assembly-based modular toolbox with different broad host range vectors by Schuster and Reisch 2021 [17]. Previously, our group carried out a small-scale characterization of inducible expression systems for *Z. mobilis* that compared TetR-P_tet_, LacI-P_lacT7A1_O3O4_ and XylS-P_m_ [11]. This work builds on top of this study and further extends it. Relevant regulator candidates are NahR, a LysR family activator originating from a plasmid found in *Pseudomonas putida* [18], LuxR an activator originating from *Vibrio fischeri* [19], VanR a GntR family repressor originating from *Caulobacter crescentus* [20] and CinR a LuxR family activator originating from *Rhizobium leguminosarum* [21]. All of these have been optimized for *E. coli* by Meyer et al. 2019 [16]. It should be briefly mentioned, that XylS is an AraC family activator originating from *Pseudomonas putida* [17], TetR is a repressor from an *E. coli* mobile element [22] and the original LacI, a repressor from *E. coli* [23]. The commonly used inducers for all of these regulators diffuse through the membrane of *E. coli* and are enriched inside the cell [16,17]. How exactly they behave in regard to the high hopanoid content membrane of *Z. mobilis* is speculative.

## Materials and Methods

### Strains and Media

For cloning of all plasmids *E. coli* NEB5α (New England Biolabs) was used. Cultivation was carried out in LB_0_ medium (10 g/L tryptone, 5 g/L yeast extract, 5 g/L NaCl) at 37 °C. For the plasmid characterization *Z. mobilis* strain ZM4 (ATCC 31821) was used and cultivated in ZM (complex) medium (bacto peptone 10 g/L, yeast extract 10 g/L, glucose 20 g/L; DSMZ GmbH) at 30 °C. For *E. coli* strains bearing plasmid, 30 µg/mL kanamycin was used. For *Z. mobilis* strains bearing plasmid, 100 µg/mL kanamycin was used. For conjugation of plasmids into ZM4 the *E. coli* strain ST18 (*pro thi hsdR*^+^ Tp^r^ Sm^r^; chromosome::RP4-2 Tc::Mu-Kan::Tn7 λ*pir* Δ*hemA*, DSM 22074) [24] was used as a donor. Conjugation was carried out after Behrendt et al. 2024 [31]. For information on inducers see supplemental Table S1.

### Plasmids

For the construction of all plasmids the Zymo-Parts assembly framework was followed [11]. For detailed instructions of the cloning procedure, see Behrendt et al. 2022 [11]. A graphical overview of the assembly of level 1 expression constructs can be found in supplemental Figure S1. Furthermore, all plasmid sequences relevant to this work can be found in the Edmond repository as annotated GeneBank files (https://doi.org/10.17617/3.F74G5A).

### Fluorescence screening

Strains of ZM4 with the respective plasmid were streaked out after conjugation, to get proper single colonies. Single colonies were picked, inoculated into ZM medium and grown overnight. The next morning the overnight cultures were diluted (1/20 v/v) into fresh ZM medium containing the respective inducer concentration. Of each culture 100 µL were directly transferred into a single well of a 96-well plate (Costar, Corning Inc.), the plate was sealed with a gas-permeable membrane (Breathe-Easy, DiversifiedBiotech). Cultivation was carried out in a VANTAstar microplate reader system (BMG LABTECH GmbH) at 100 rpm, to prevent cell agglomeration and provide oxygen for maturation of mCherry. Biomass was measured via absorbance at 600 nm wavelength (OD_600_), at the center of each well with volume-based pathlength correction. mCherry fluorescence was measured with excitation at 570 nm (15 nm bandwidth) and emission at 620 nm (20 nm bandwidth), using 20 flashes per well, a settling time of 0.5 s, and enhanced dynamic range. OD600 and fluorescence were recorded every 30 min. Raw measurements were blank-corrected using wells containing ZM medium only. Seven independent cultivations were performed.

### Data analysis

The cultivation data were analyzed and plotted using R. The data frame created from the experiments and used for the plots, as well as the entire R code needed to generate the Hill slope and all the plots can be found in the Edmond repository (https://doi.org/10.17617/3.F74G5A).

## Results and discussion

In this work NahR-P_salTTC_, VanR^AM^-P_vanCC_, CinR^AM^-P_cin_ and LuxR-P_luxB_ were tested for application in *Z. mobilis* for the first time and compared to the commonly used inducible expression systems TetR-P_tet_ and LacI-P_lacT7A1_O3O4_ and also XylS-P_m_. A general schematic of the genetic basis of the elements characterized in this work can be found in Figure 1.

**Figure 1.**
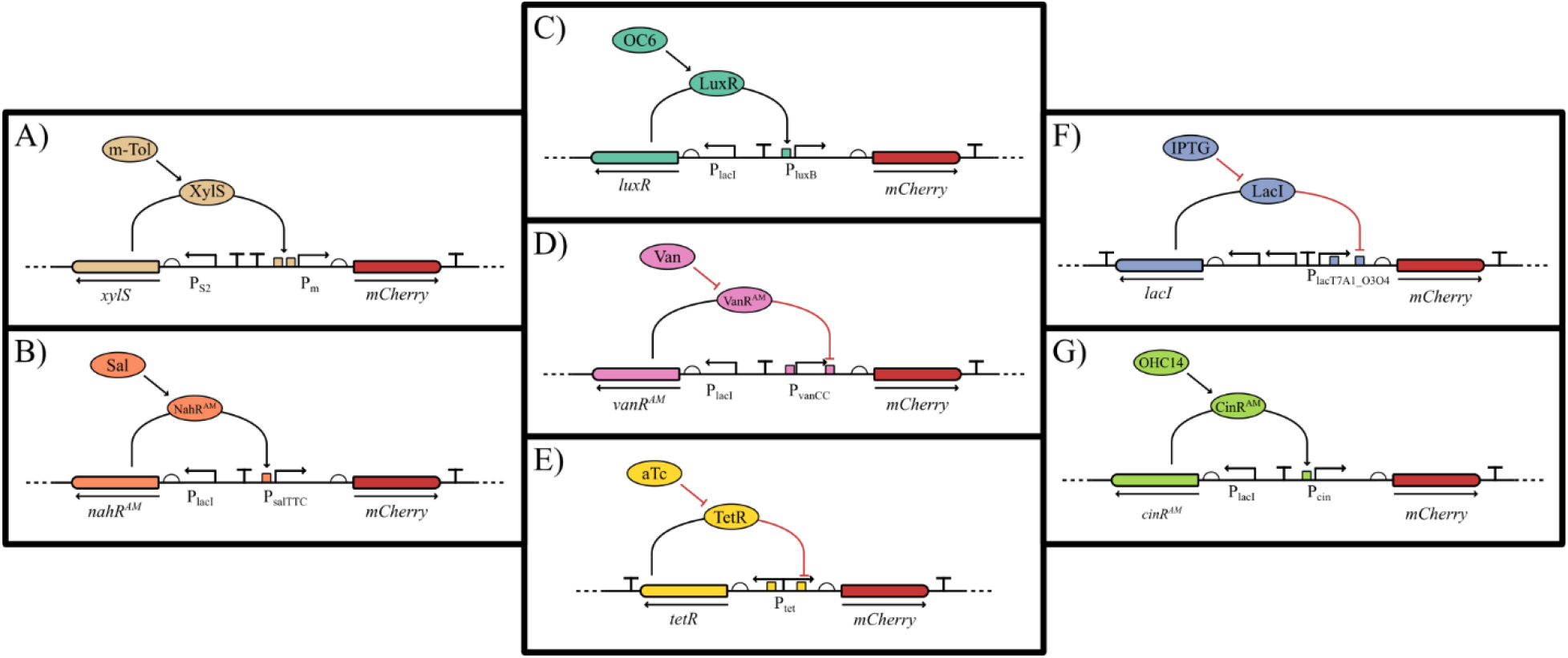
Graphic overview of the transcription regulators investigated in this work. In all systems addition of the chemical inducer increases transcription. Whether activation occurs through inhibition of a repressor or activation of an activator is shown. Except for the regulator-promoter region, all genetic constructs characterized in this study have the same genetic background. A) shows induction via m-toluate (m-Tol), activating XylS, which binds to a region upstream of P_m_. B) shows the induction via salicylic acid (Sal), which activates NahR^AM^, which binds to a region upstream of P_salTTC_. C) shows induction via N-(3-oxohexanoyl) homoserine lactone (OC6), which binds to LuxR, which binds to a region upstream of P_luxB_. D) shows the induction via vanillic acid (Van), which binds to VanR^AM^, which binds to regions flanking P_vanCC_. E) shows the induction via anhydrotetracyclin, (aTc) which binds to TetR, which binds to regions inside the bidirectional P_tet_. F) shows the induction via isopropyl-β-d-thiogalactopyranosid (IPTG), which binds to LacI, which binds to regions inside the space of and downstream of P_lacT7A1_O3O4_. G) shows the induction via hydroxytetradecanoyl-homoserine lactone (OHC14), which binds to CinR^AM^, which binds to a region upstream of P_cin_. Color coding of the regulator-promoter pairs is resumed in the rest of this manuscript. The genetic circuits presented here are aligned with Synthetic Biology Open Language (SBOL).

To test these genetic regulators in a standardized background, they were adapted to the modular cloning framework Zymo-Parts [11]. Therefore, plasmid construction was carried out with Golden Gate cloning [27], a cloning schematic can be found in supplemental Figure S1. References for the regulator-promoter pairs can be found in [11,16,28,29]. All inducible transcription systems have been paired with the same strong rbs (rbs10k, [11,30]), *mcherry* as a fluorescence reporter gene [11,31] and a terminator (T_soxR_ [11,32]). The backbone of the resulting constructs contains a kanamycin resistance cassette, the pUC ori for high-copy replication in *E. coli* and the pZMO7 replication apparatus for stable and homogeneous replication in *Z. mobilis* [25,26]. TetR-P_tet_, LacI-P_lacT7A1_O3O4_ and XylS-P_m_ have previously been analyzed via flow cytometry in our group [11], however, with a different genetic background and only at a single time point of cultivation. In this work, the whole characterization was carried out in the VANTAstar microplate reader system (BMG LABTECH GmbH). Comparisons of the results generated in this work are made to a previous study from our group [11], the characterization done in *E. coli* by Meyer et al. 2019 [16], the plasmid toolbox for controlled gene expression across the Proteobacteria from Schuster and Reisch 2021 [28] and a recent investigation of inducible transcription systems in *Rhodobacter sphaeroides* [29]. The comparison to *R. sphaeroides* is made as it is also an Alphaproteobacterium and the underlying cloning framework used in the reference is the same, which is used in this work. As references for the inducible transcription regulation, five constitutive promoters, that were previously characterized for *Z. mobilis* [11], were used with the same genetic background as the inducible systems. These constitutive expression systems are three strong native promoters of *Z. mobilis* [11,30] and two synthetic promoters designed with the promoter calculator [33]: P_gap_, P_pdc334_, P_tuf_, P_strong1k_ and P_strong100k*_. A minimum of 8 and a maximum of 14 different inducer concentrations were tested for each promoter (see supplemental Table S2). Growth was only slightly inhibited by the addition of vanillic acid at the highest concentrations; otherwise, the inducers did not interfere with growth at the tested concentrations (see supplemental Figure S2).

The dynamic range of fluorescence observed for each inducible transcription system over time can be found in Figure 2. Ten hours after inoculation generally marks the transition from exponential growth to stationary phase in the experimental setup used in this study (see supplemental Figure S2). For all tested inducible transcription systems, the ten-hour mark was the point of lowest basal reporter expression (see supplemental Figures S3 and S4). Afterwards, during stationary phase, the fluorescence normalized by biomass is slowly but steadily increasing, to varying degrees for the different regulation systems. The most likely explanation is that full maturation of mCherry is delayed during growth, as it is dependent on oxygen [34] and *Z. mobilis* respiration rate should be higher during growth. This effect is not observed for the Marionette strains of *E. coli* which relied on YFP as a fluorescence reporter [16], with YFP being less dependent on oxygen for maturation [35]. Data of the first 2 h after inoculation is to be viewed with caution as the media used for correction had rather high background fluorescence in the setup used. Thus, low biomass concentrations have a higher standard deviation for mCherry fluorescence (see supplemental Figure S3).

**Figure 2.**
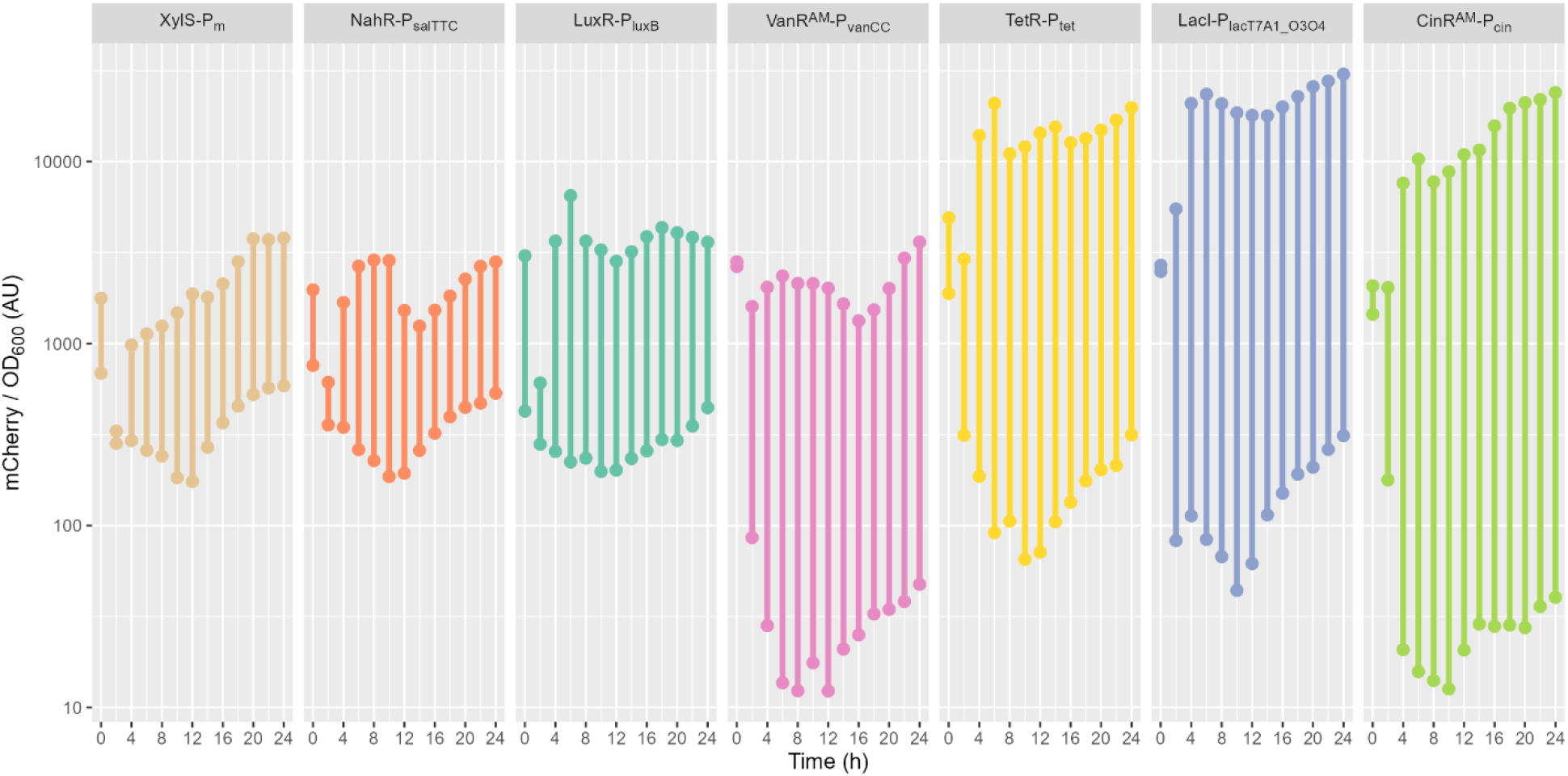
Dynamic range of mCherry fluorescence over time for each inducible transcription system. The range from lowest induction to highest induction was plotted in 2 h steps from 0 to 24 h of cultivation. For a simplified overview, see supplemental Figure S5.

Crosstalk was not assessed here. In *E. coli*, Meyer et al. 2019 [16] reported minimal crosstalk for these systems, with the exception of m-toluate which was not part of their study and aTc which showed low crosstalk with P_tac_. Whether this holds in *Z. mobilis* remains to be tested.

Titration curves with Hill fits can be found in Figure 3, the ten-hour mark of cultivation was used for this plot. Characteristics of the Hill fit (Hill coefficient, EC_50_, dynamic range of expression, as well as basal and maximum expression as a multiple of the expression generated by P_strong1k_) can be found in Table 1. An expanded table showing these characteristics for the same regulatory systems in other hosts can be found in supplemental Table S3. The exact numbers can be found in the tables; in the text approximations indicated as “about” are used for readability.

**Table 1:** Details of the titration results for each inducible transcription system. The Hill equation describes the binding of ligands (inducers) to macromolecules (regulator proteins). The Hill coefficient (*n*) describes cooperativity of ligand binding and thus the steepness of the Hill slope. n > 1 describes a positively cooperative binding of ligands and a steep slope, while n = 1 describes non-cooperative binding and n < 1 describes negatively cooperative binding of ligands and a gentle slope. The EC_50_ is the inducer concentration, at which 50% of the maximal response is reached. The dynamic range describes the fold between minimal/basal (y_min_) and maximal (y_max_) expression. Minimal and maximal expression themselves are shown as multiples of the constitutive expression derived from the weak promoter P_strong1k_. For the Hill coefficient and EC_50_ standard error are shown as well, for dynamic range, y_min_ / P_strong1k_ and y_max_ / P_strong1k_ approximate standard errors are shown, for more details see the R code.

| System | Hill coefficient | $EC_{50}$ [μM] | Dynamic range (fold) | $y_{min} / y_{Pstrong1k}$ | $y_{max} / y_{Pstrong1k}$ |
| --- | --- | --- | --- | --- | --- |
| XylS-$P_m$ | $1.1 \pm 0.31$ | $238 \pm 1.2e+02$ | $8.06 \pm 1.7$ | $1.99 \pm 0.49$ | $16 \pm 3$ |
| NahR-$P_{salTTC}$ | $0.75 \pm 0.34$ | $1.69e+06 \pm 1e+09$ | $15.4 \pm 3.5$ | $2.02 \pm 0.38$ | $31 \pm 8$ |
| LuxR-$P_{luxB}$ | $0.88 \pm 0.12$ | $37.8 \pm 16$ | $16.5 \pm 1.3$ | $2.15 \pm 0.36$ | $36 \pm 6$ |
| VanR<sup>AM</sup>-$P_{vanCC}$ | $1.23 \pm 0.13$ | $606 \pm 2.8e+02$ | $121 \pm 41$ | $0.19 \pm 0.07$ | $23 \pm 4$ |
| TetR-$P_{tet}$ | $1.88 \pm 0.1$ | $0.057 \pm 0.01$ | $185 \pm 39$ | $0.71 \pm 0.18$ | $131 \pm 22$ |
| LacI-$P_{lacT7A1\_O3O4}$ | $1.28 \pm 0.048$ | $284 \pm 55$ | $421 \pm 1.3e+02$ | $0.48 \pm 0.16$ | $201 \pm 34$ |
| CinR<sup>AM</sup>-$P_{cin}$ | $1.15 \pm 0.053$ | $1.29 \pm 0.3$ | $696 \pm 3e+02$ | $0.14 \pm 0.06$ | $96 \pm 15$ |

**Figure 3.**
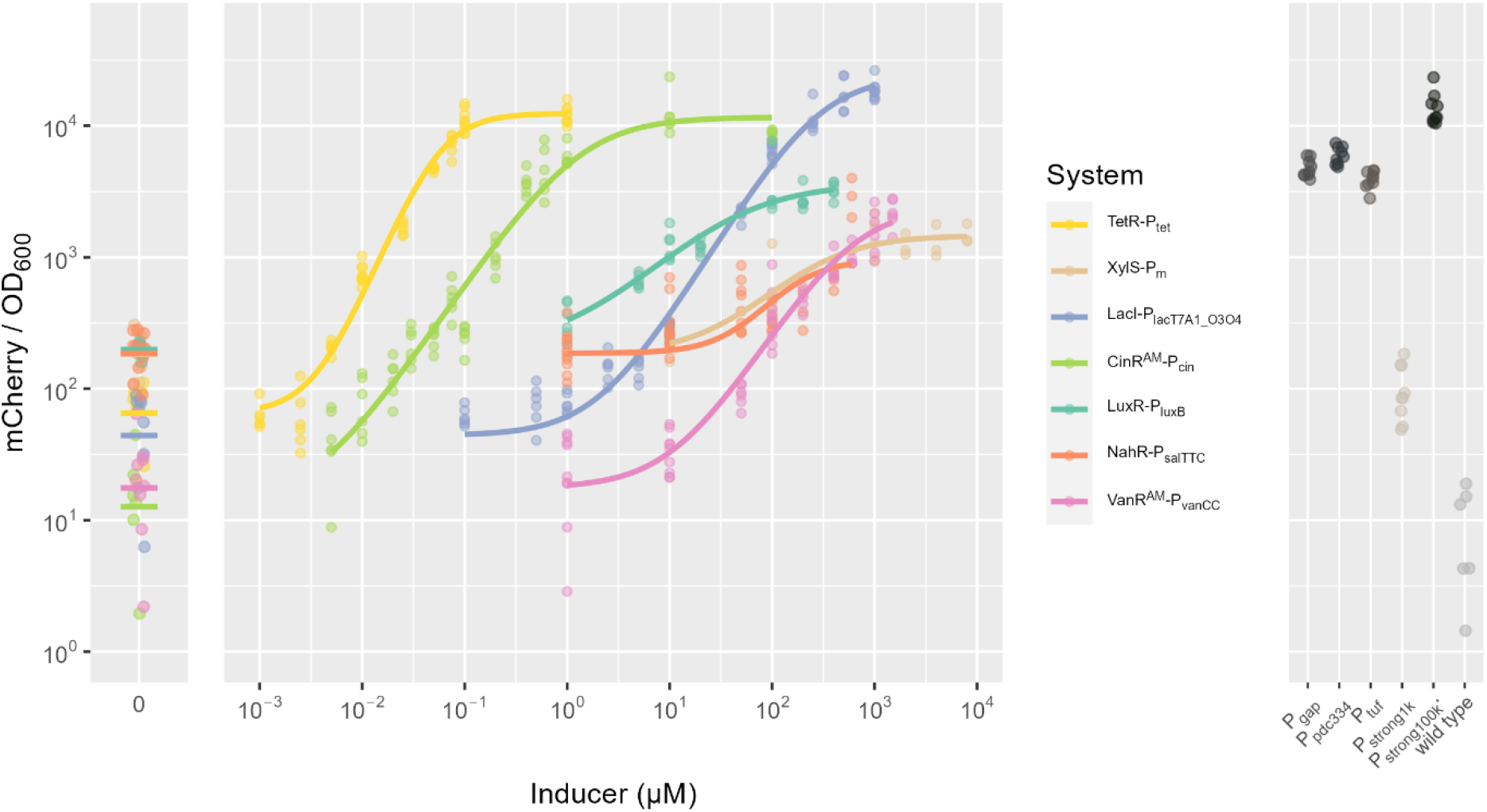
Titration curves with Hill fit and points of reference. The ten-hour mark of the cultivation was used as a reference point to generate the plot. Each replicate is shown as a single dot.

Across the seven tested inducible transcription regulation systems, CinR^AM^-P_cin_ and LacI-P_lacT7A1_O3O4_ showed the largest dynamic ranges (about 700-fold and 400-fold, respectively), indicating strong inducibility with low basal expression relative to maximal expression. TetR-P_tet_ and VanR^AM^-P_vanCC_ also displayed high dynamic ranges (about 200-fold and 120-fold, respectively). In contrast, XylS-P_m_ exhibited the smallest dynamic range (about 8-fold), reflecting limited separation between basal and induced states under the tested conditions, which aligns well with previous characterizations of the system in *Z. mobilis* [11]. NahR-P_salTTC_ and LuxR-P_luxB_ both had relatively low dynamic ranges as well (both about 15-fold). By contrast, the dynamic range of NahR-P_salTTC_ in *R. sphaeroides* was one order of magnitude greater and the dynamic range of LacI-P_lacT7A1_O3O4_ a magnitude smaller, while for VanR^AM^-P_vanCC_ the results were similar [29].

Sensitivity, quantified by EC_50_, varied widely among systems. TetR-P_tet_ and CinR^AM^-P_cin_ were the most sensitive (about 0.06 µM and 1.3 µM), whereas XylS-P_m_, LacI-PlacT7A1_O3O4, and VanR^AM^-P_vanCC_ required substantially higher inducer concentrations for half-maximal activation (EC_50_ in the 10^2^-10^3^ µM range). In between LuxR-P_luxB_ is found (about 40 µM). The NahR-P_salTTC_ fit yielded an extremely large and uncertain EC_50_ (about 1.7×10^6^ ± 10^9^ µM), suggesting that half-maximal induction was not well constrained by the assayed concentration range or rather that the response did not reach a clear saturation regime.

To benchmark performance against a constitutive reference, basal and maximal expression were normalized to weak, constitutive, synthetic promoter P_strong1k_ (Table 1). LacI-P_lacT7A1_O3O4_, TetR-P_tet_ and CinR^AM^-P_cin_ reached the highest maximal expression relative to P_strong1k_ (about 200, 130 and 95 of P_strong1k_, respectively), exceeding the expression levels of the strong native promoters (P_gap_, P_pdc334_ and P_tuf_), with LacI-P_lacT7A1_O3O4_ even reaching levels of the strongest known constitutive promoter for *Z. mobilis* (P_strong100k*_). Notably, CinR^AM^-P_cin_ also had the lowest basal expression (about 0.1 of P_strong1k_). For VanR^AM^-P_vanCC_ the second lowest basal expressions (about 0.2 of P_strong1k_) was observed, however, the maximal induction was rather low (max about 23 of P_strong1k_). In contrast, XylS-P_m_, NahR-P_salTTC_ and LuxR-P_luxB_ all showed rather high levels of basal expression (about 2 of P_strong1k_), rendering them leaky.

As the version of CinR^AM^-P_cin_, VanR^AM^-P_vanCC_, NahR-P_salTTC_ and LuxR-P_luxB_ characterized here all were optimized for *E. coli*, there will be no attempts to map phylogenetic relatedness of the original host (e.g., VanR from *C. crescentus*) to functionality in *Z. mobilis*. However, the LuxR-P_luxB_ and NahR-P_salTTC_ transcriptional regulation systems tested here also demonstrated comparably low dynamic ranges when tested in other Alphaproteobacteria (*Agrobacterium fabrum* C58, *Sulfitobacter* sp. EE-36 and *Ruegeria* sp. TM1040), while the CinR^AM^-P_cin_ pair demonstrated high dynamic range in those hosts [28].

Overall, the dataset demonstrates that inducible systems differ markedly in both operational range (EC_50_ and slope) and achievable expression window (basal leakage and maximal output), enabling selection of regulators tailored to either low-leakage control (VanR^AM^-P_vanCC_ and CinR^AM^-P_cin_), high maximal expression (LacI-P_lacT7A1_O3O4_, CinR^AM^-P_cin_ and TetR-P_tet_), or high sensitivity (TetR-P_tet_ and CinR^AM^-P_cin_). Our group previously switched the native P_pdc_ promoter on the chromosome of *Z. mobilis* for the inducible LacI-P_lacT7A1_O3O4_ system to channel metabolic flux away from ethanol production [36], with the new results shown here, VanR^AM^-P_vanCC_ and CinR^AM^-P_cin_ might have been better choices, as their basal expression is lower. Especially, VanR^AM^-P_vanCC_ with vanillic acid as a cheap inducer would have been an interesting candidate, regarding commercial viability of bulk chemical production.

Finally, defining an optimal heterologous transcription regulator as having background expression (uninduced) lower than the constitutive expression of the weakest constitutive promoter assessed before (P_strong1k_, [11]) and a dynamic expression range of more than 100-fold, this holds true for four of the systems tested here: VanR^AM^-P_vanCC_, TetR-P_tet_, LacI-P_lacT7A1_O3O4_ and CinR^AM^-P_cin_.

## Conclusion

All regulator-promoter pairs assessed in this study demonstrate functionality in *Z. mobilis*, however, to different extents. The activator-promoter pairs XylS-P_m_, NahR-P_salTTC_ and LuxR-P_luxB_ appear unsuitable for most applications in *Z. mobilis*, as they all share a high basal expression and low dynamic range. In contrast, VanR^AM^-P_vanCC_, TetR-P_tet_, LacI-P_lacT7A1_O3O4_ and CinR^AM^-P_cin_ all offer characteristics suitable for engineering applications and investigation of its physiology and metabolism. Specifically, the newly added regulator-promoter pairs VanR^AM^-P_vanCC_ and CinR^AM^-P_cin_ offer very low basal expression, outperforming the already established inducible systems and in the case of VanR^AM^-P_vanCC_ also having a very cost-effective inducer in the form of vanillic acid.

## Author contributions

NA

## Acknowledgements

ChatGPT-5.2 (OpenAI) assisted with the R code and its annotation.

The author gratefully acknowledges the funding by the European Regional Development Fund (ERDF) within the programme Research and Innovation (FerMeVi project).

## Declaration of competing interest

The author declares no competing interest

## Additional information

Supporting information file 1: Contains supporting figures and tables.

Sequences of the plasmids, the data frame with cultivation data and annotated R code are available at Edmond: https://doi.org/10.17617/3.F74G5A

## Notes

https://doi.org/10.17617/3.F74G5A

### Competing Interest Statement

The authors have declared no competing interest.

