## Supporting information file 1 for "Seven inducible promoters for *Zymomonas mobilis*"

Gerrich Behrendt*

Analysis and Redesign of Biological Networks, Max Planck Institute for Dynamics of Complex Technical Systems, 39106 Magdeburg, Germany

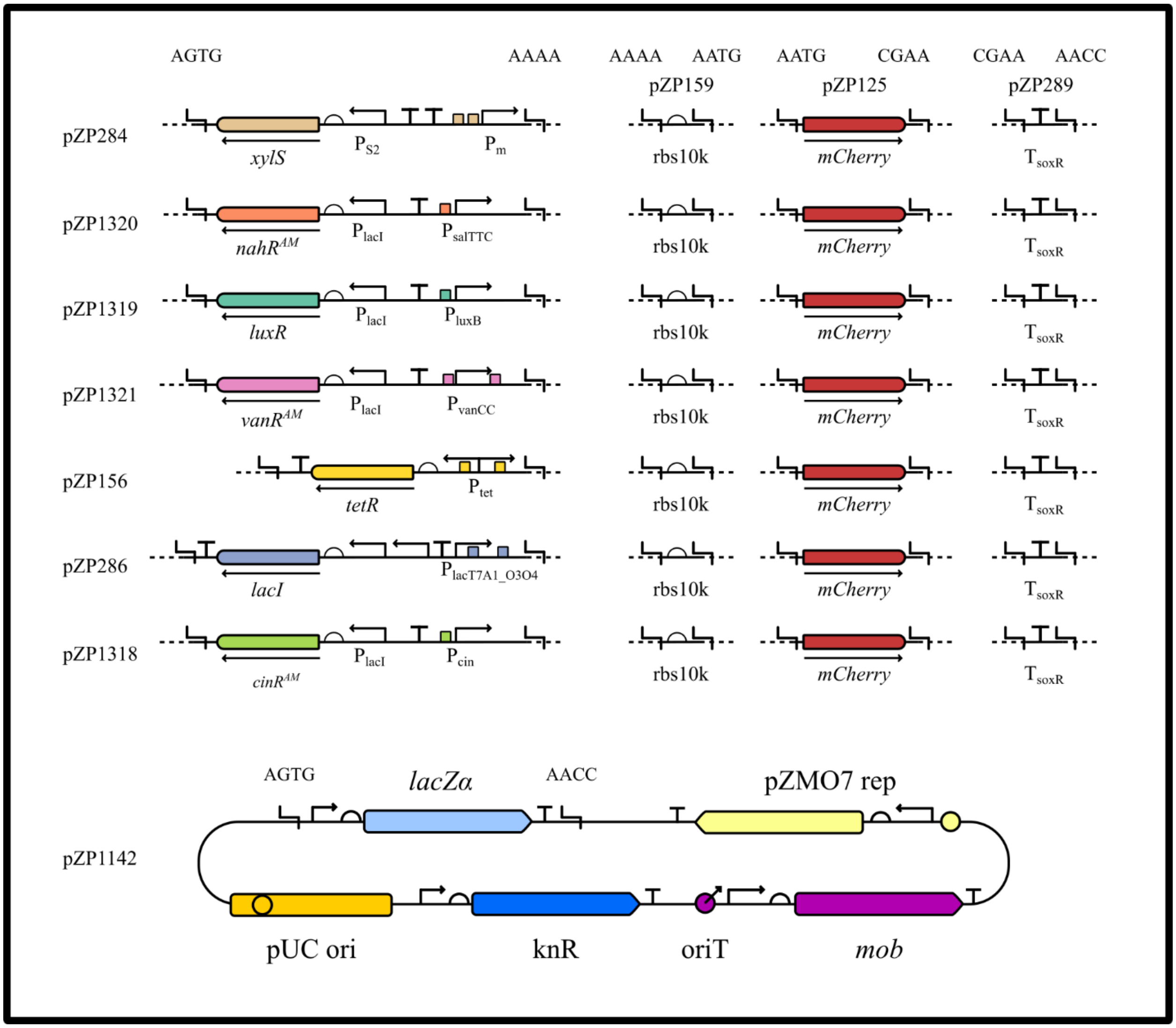


Supplemental Figure S1: Graphical cloning schematic of assembly parts used in this study. SBOL standardized glyphs are used.

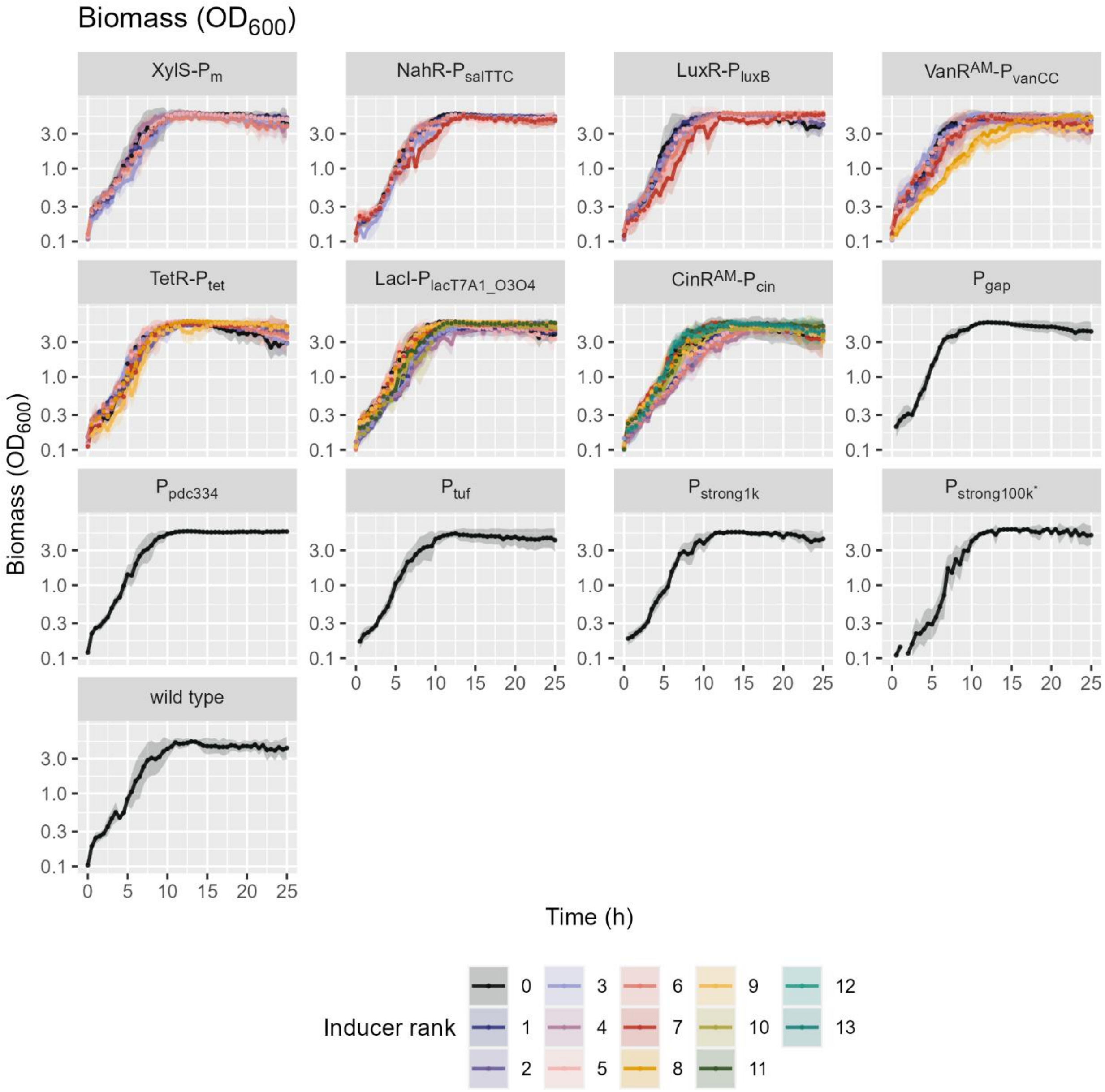


Supplemental Figure S2: Growth of all VANTAstar cultivations carried out in this study. Shown are two-sided 95% confidence intervals with the geometric mean. Inducers are only labeled by rank to obscure the different concentration ranges and instead indicate and increase or decrease of concentration for each compound. See Table S1 for reference of inducer concentration.

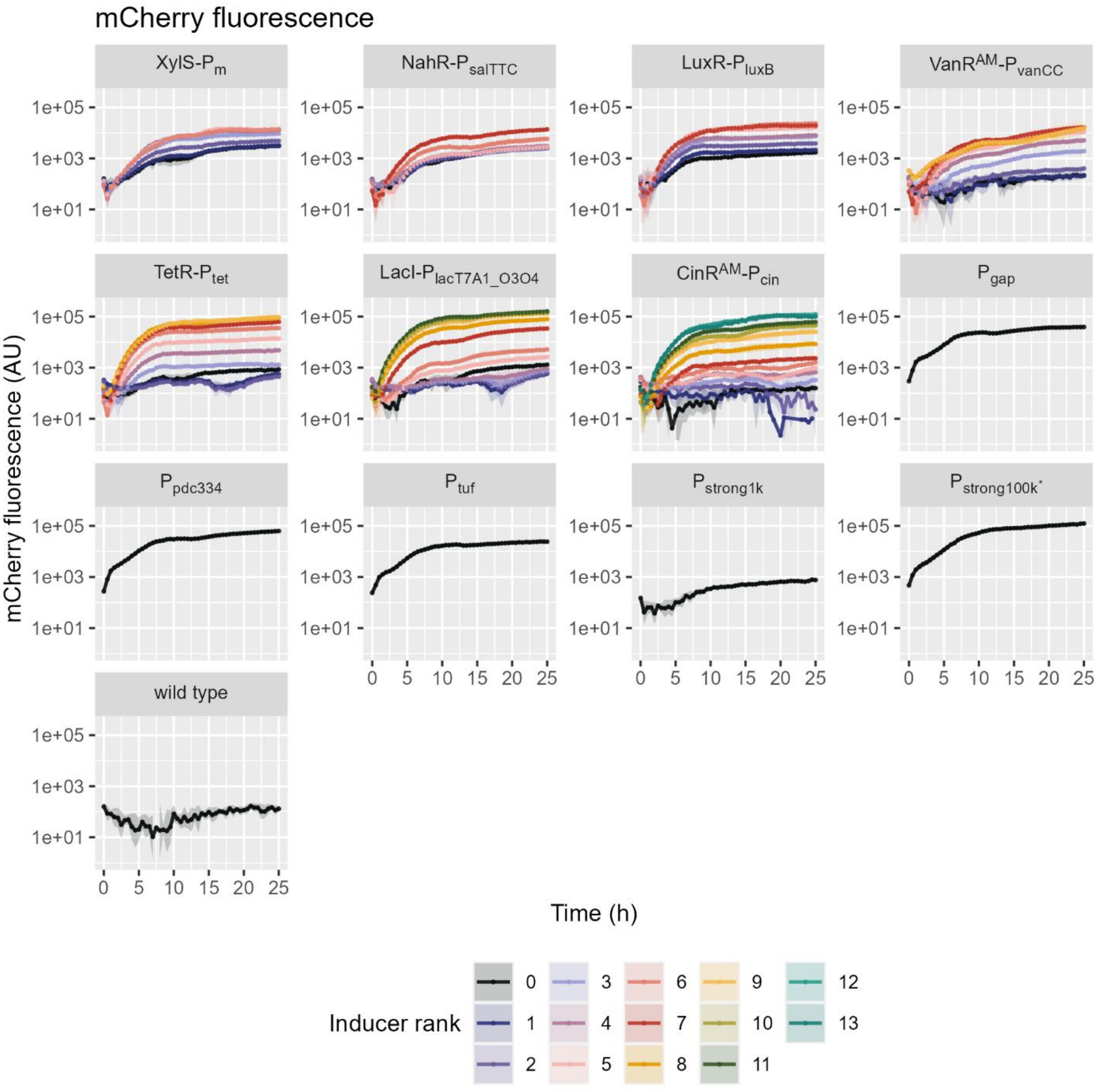


Supplemental Figure S3: mCherry fluorescence of all VANTAstar cultivations carried out in this study. Shown are two-sided 95% confidence intervals with the geometric mean. Inducers are only labeled by rank to obscure the different concentration ranges and instead indicate and increase or decrease of concentration for each compound. See Table S1 for reference of inducer concentration.

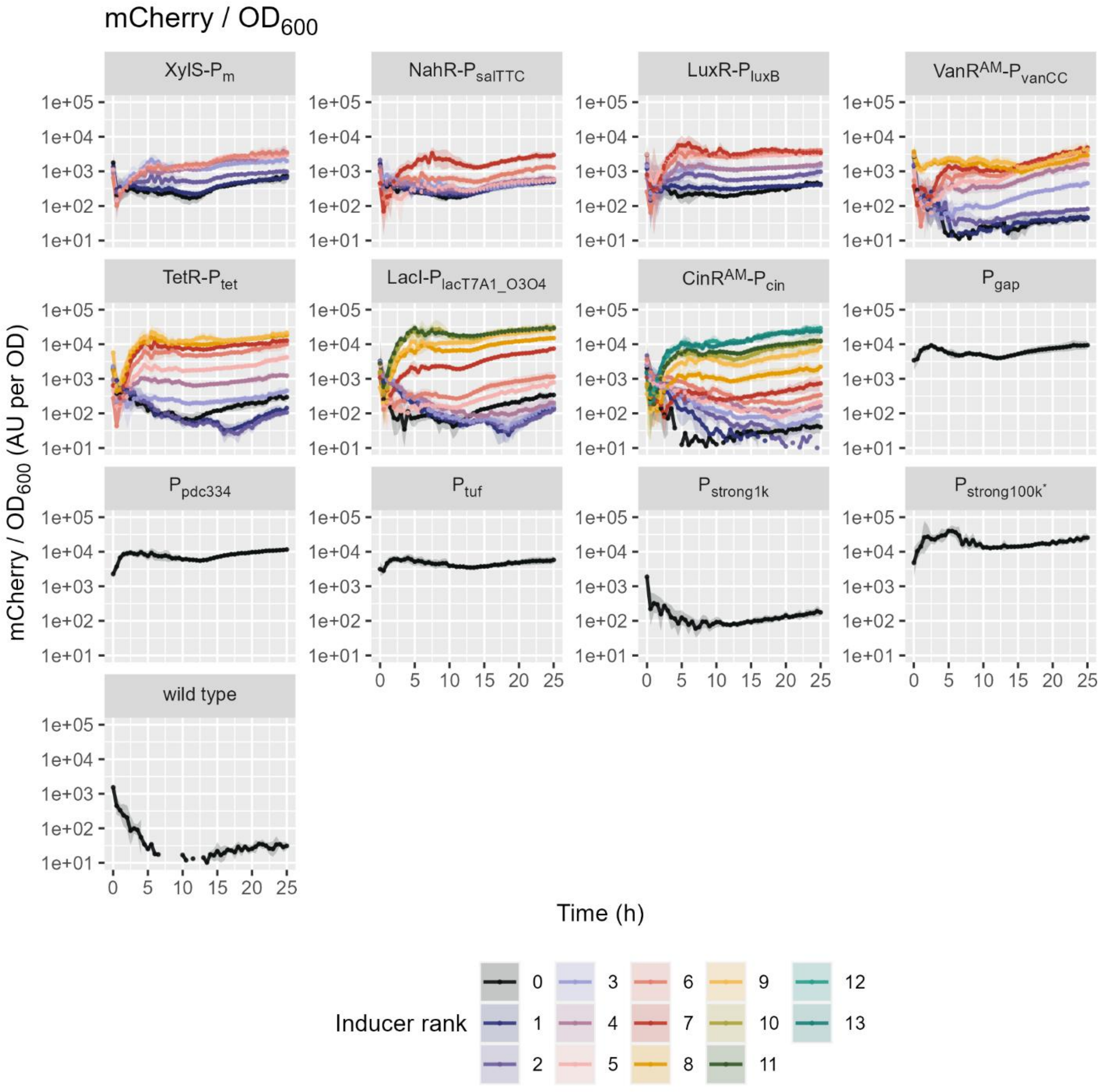


Supplemental Figure S4: mCherry fluorescence normalized to $OD_{600}$ of all VANTAstar cultivations carried out in this study. Shown are two-sided 95% confidence intervals with the geometric mean. Inducers are only labeled by rank to obscure the different concentration ranges and instead indicate and increase or decrease of concentration for each compound. See Table S1 for reference of inducer concentration.

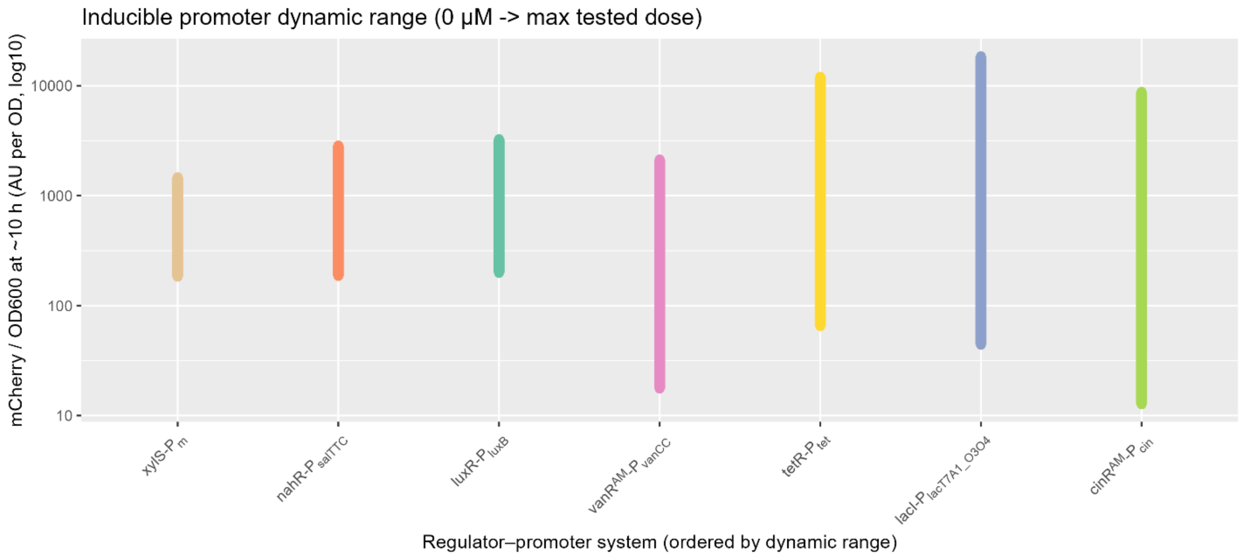


Supplemental Figure S5: Dynamic range of mCherry fluorescence over time for each inducible transcription system at the 10.

Supplemental Table S1: Substances used as inducers for the respective transcriptional regulators. Stocks were made either at 1000-times the applied inducer concentration (see supplemental Table S2) or the maximal solubility.

| **System** | **Inducer** | **Source** | **Solvent** |
|---|---|---|---|
| XylS-$P_{m}$ | m-toluic acid (m-Tol) | Sigma, T36609 | Dimethyl sulfoxide (DMSO) |
| NahR-$P_{salTTC}$ | Salicylic acid (Sal) | Sigma, 247588 | Water |
| LuxR-$P_{luxB}$ | N-(3-oxohexanoyl) homoserine lactone (OC6) | Sigma, 51481 | Dimethyl sulfoxide (DMSO) |
| VanR$^{AM}$-$P_{vanCC}$ | Vanillic acid (Van) | Sigma, H36001 | Ethanol |
| TetR-$P_{tet}$ | Anhydrotetracyclin (aTc) | Sigma, 37919 | Water |
| LacI-$P_{lacT7A1\_O3O4}$ | Isopropyl-ß-D-thiogalactopyranosid (IPTG) | Carl ROTH, CN08 | Water |
| CinR$^{AM}$-$P_{cin}$ | Hydroxytetradecanoyl-homoserine lactone (OHC14) | Sigma, K3007 | Dimethyl sulfoxide (DMSO) |

Supplemental Table S2:Numbered inducer concentrations tested for each of the transcription regulators as µM. Meant to help the readability of Figures S2, S3 and S4.

| **Plasmid** | **Ind_0** | **Ind_1** | **Ind_2** | **Ind_3** | **Ind_4** | **Ind_5** | **Ind_6** | **Ind_7** | **Ind_8** | **Ind_9** | **Ind_10** | **Ind_11** | **Ind_12** | **Ind_13** |
|---|---|---|---|---|---|---|---|---|---|---|---|---|---|---|
| XylS-$P_{m}$ | 0 | 10 | 100 | 1000 | 2000 | 4000 | 8000 | 16000 | NA | NA | NA | NA | NA | NA |
| NahR-$P_{salTTC}$ | 0 | 1 | 10 | 50 | 100 | 200 | 400 | 600 | NA | NA | NA | NA | NA | NA |
| LuxR-$P_{luxB}$ | 0 | 1 | 5 | 10 | 20 | 100 | 200 | 400 | NA | NA | NA | NA | NA | NA |
| VanR$^{AM}$-$P_{vanCC}$ | 0 | 1 | 10 | 50 | 100 | 200 | 400 | 600 | 1000 | 1500 | NA | NA | NA | NA |
| TetR-$P_{tet}$ | 0 | 0.001 | 0.003 | 0.005 | 0.01 | 0.025 | 0.05 | 0.075 | 0.1 | 1 | NA | NA | NA | NA |
| LacI-$P_{lacT7A1\_O3O4}$ | 0 | 5 | 10 | 50 | 100 | 250 | 500 | 1000 | 0.1 | 0.5 | 1 | 2.5 | NA | NA |
| CinR$^{AM}$-$P_{cin}$ | 0 | 0.005 | 0.01 | 0.02 | 0.03 | 0.05 | 0.075 | 0.1 | 0.2 | 0.4 | 0.6 | 1 | 10 | 100 |

Supplemental Table S3: Details of the titrations for all inducible transcription systems tested in this work, expanded with results from Meyer et al. 2019 for *E. coli* DH10B (blue) and Kostanjšek et al. 2026 for *R. sphaeroides* (red). The dynamic range for *E. coli* DH10B was calculated from the $y_{min}$ $y_{max}$ values for fluorescence provided by Meyer et al. 2019. The $EC_{50}$ value is described as K in Meyer et al. 2019.

| **System** | **Hill slope** | **$EC_{50}$ (/K) [µM]** | **Dynamic range [fold]** | **Basal / $P_{strong1k}$** | **Max / $P_{strong1k}$** |
|---|---|---|---|---|---|
| XylS-$P_m$ | 1.1 ± 0.31 | 238 ± 1.2e+02 | 8.06 ± 1.7 | 1.99 ± 0.49 | 16 ± 3 |
| NahR-$P_{salTTC}$ | 0.75 ± 0.34 | 1.69e+06 ± 1e+09 | 15.4 ± 3.5 | 2.02 ± 0.38 | 31 ± 8 |
| NahR-$P_{salTTC}$ *E. coli* DH10B | 1.8 | 43 | 595 | | |
| NahR-$P_{salTTC}$ *R. sphaeroides* | 1.89 ± 0.22 | 8.8 ± 0.5 | 136 ± 5.6 | | |
| LuxR-$P_{luxB}$ | 0.88 ± 0.12 | 37.8 ± 16 | 16.5 ± 1.3 | 2.15 ± 0.36 | 36 ± 6 |
| LuxR-$P_{luxB}$ *E. coli* DH10B | 1.8 | 0.12 | 542 | | |
| VanR$^{AM}$-$P_{vanCC}$ | 1.23 ± 0.13 | 606 ± 2.8e+02 | 121 ± 41 | 0.19 ± 0.07 | 23 ± 4 |
| VanR$^{AM}$-$P_{vanCC}$ *E. coli* DH10B | 2.3 | 26 | 1250 | | |
| VanR$^{AM}$-$P_{vanCC}$ *R. sphaeroides* | 0.91 ± 0.1 | 310 ± 60 | 432 ± 192 | | |
| TetR-$P_{tet}$ | 1.88 ± 0.1 | 0.057 ± 0.0074 | 185 ± 39 | 0.71 ± 0.18 | 131 ± 22 |
| LacI-$P_{lacT7A1\_O3O4}$ | 1.28 ± 0.048 | 284 ± 55 | 421 ± 1.3e+02 | 0.48 ± 0.16 | 201 ± 34 |
| LacI-$P_{lacT7A1\_O3O4}$ *R. sphaeroides* | 1.55 ± 0.22 | 59 ± 7 | 27 ± 2 | | |
| CinR$^{AM}$-$P_{cin}$ | 1.15 ± 0.053 | 1.29 ± 0.3 | 696 ± 3e+02 | 0.14 ± 0.06 | 96 ± 15 |
| CinR$^{AM}$-$P_{cin}$ *E. coli* DH10B | 2.3 | 0.43 | 500 | | |
